# Cell-state mapping reveals a reversible neuroblast accumulation in the aging mouse hippocampus

**DOI:** 10.64898/2026.04.23.717949

**Authors:** Natalí B. Rasetto, Magalí Herrero, Ariel A. Berardino, Daniela J. Di Bella, Damiana Giacomini, Mariela F. Trinchero, Paola Arlotta, Ariel Chernomoretz, Alejandro F. Schinder

## Abstract

Hippocampal neurogenesis supports learning and memory by generating new granule cells throughout life. However, both the rate and speed of this process have been shown to decline with age in mice. Currently, the molecular determinants underlying the effects of aging on neuronal development and differentiation remain unknown. To elucidate these mechanisms, we combined lineage tracing with single-nucleus RNA sequencing of adult-born granule cells (aGCs) in the aging mouse hippocampus. Employing this temporally-resolved transcriptional atlas of aGCs development, we observed that aging leads to a marked accumulation of postmitotic neuroblasts (NBs), revealing a stage-specific bottleneck in the neurogenic trajectory. Voluntary running reduced NB accumulation by restoring the progression through this developmental impasse, allowing neuronal maturation to be resumed. This shift was accompanied by a significant reduction in apoptosis at the NB stage. Together, these findings highlight postmitotic NBs as a developmental stage of increased vulnerability during aging and point to the restorative effects of exercise in reducing apoptosis and driving cellular transitions towards a neurogenic trajectory.

## INTRODUCTION

The hippocampus is critical for the formation, consolidation, and retrieval of memories, and plays a key role in generating and maintaining spatial representations (1, 2). Within this structure, the dentate gyrus (DG) is notable for its capacity to generate new neurons throughout life. In young mice, adult-born granule cells (aGCs) progress through well-defined developmental stages and require over ten weeks to reach full maturity and functionally integrate into existing circuits (3–5). Aging is associated with a significant reduction in neurogenesis, reduced global electrical activity in the DG, and a substantial delay in the maturation and functional integration of newly generated neurons, collectively contributing to the impairment of brain plasticity observed in this condition (6–9). The decline in neurogenesis is largely driven by an increased quiescence of radial glia-like cells (RGL), altered proteostasis, and reduced asymmetric divisions, contributing to stem cell pool depletion and a shift toward gliogenesis (10–13). In parallel, newly generated neurons exhibit lower survival rates, partly due to increased apoptosis and diminished trophic and neurotransmitter support (13–16). Moreover, neurons born in middle aged animals already exhibit a protracted development. In fact, three-week-old neurons display morphological and functional properties typical of one-week-old neurons in young animals (9). Nevertheless, environmental stimuli such as physical exercise or enriched environment can partially rescue neurogenesis and functional integration in the aged brain, indicating that key components of neurogenic plasticity remain intact despite aging (9, 17, 18). Interestingly, these interventions also improve learning and memory performance (19–23).

Recent advances in single-cell and spatial transcriptomic technologies have enabled comprehensive profiling of the aged hippocampus under various conditions, including models of neurodegenerative diseases such as Alzheimer’s (24, 25). These studies have uncovered key molecular signatures of aging and highlighted the beneficial effects of interventions like physical exercise on the brain. However, the transcriptional dynamics of adult hippocampal neurogenesis in the aging brain remain largely unexplored. To address this gap, we employed permanent fluorescent labeling at the time of cell birth to isolate and transcriptionally profile adult-born neurons of defined ages, capturing developmental transitions that are not readily distinguishable by morphological or electrophysiological analysis. Building on our previously published single-nucleus RNA sequencing (snRNA-seq) dataset from young mice (26), we now generated a new dataset from developing aGCs in the aged hippocampus, enabling a direct comparison of transcriptional trajectories across age.

This work reveals that the core transcriptional features of aGC developmental stages are preserved with age. However, neurons accumulate at a postmitotic neuroblast (NB) stage in the aged hippocampus, indicating a discrete developmental bottleneck underlying delayed maturation. Moreover, we show that accumulated NBs are prone to suffer apoptosis. Nonetheless, aGCs from running mice showed a restored transcriptional progression beyond this stage, an increase in neuronal maturation and substantially reduced cell death. These results indicate that physical activity preserves early-stage neuron survival and enables stalled neurons to re-enter the differentiation trajectory.

Our findings provide a high-resolution and stage-specific transcriptomic map of neurogenesis in the aged hippocampus, revealing a developmental stage at which neuronal progression is disrupted in aging. Furthermore, we identify a key transcriptional checkpoint that could be targeted to restore neuronal development in aging and disease.

## RESULTS

### Profiling neurogenesis in the aging hippocampus

The mechanisms underlying the delayed aGC maturation in the aging hippocampus remain unclear. To understand the transcriptional programs orchestrating the pathway from a RGL to a mature aGC in the aging mouse, we performed a comprehensive single-nucleus RNA sequencing (snRNA-seq) analysis of aGCs obtained from 8-to 9-month-old mice (8M, aging dataset). We employed a double transgenic mouse model, *Ascl1^CreERT2^;CAG^floxStopSun1sfGFP^* that allows permanent nuclear membrane labeling of Ascl1-expressing RGLs via tamoxifen (TAM)-induced expression of Sun1-sfGFP (**Fig. 1A**) (26–28). GFP^+^ nuclei from 3-, 5-, and 8-week-old aGCs (cohorts w3, w5 and w8) were isolated by fluorescent-activated cell sorting (FACS).

**Figure 1.**
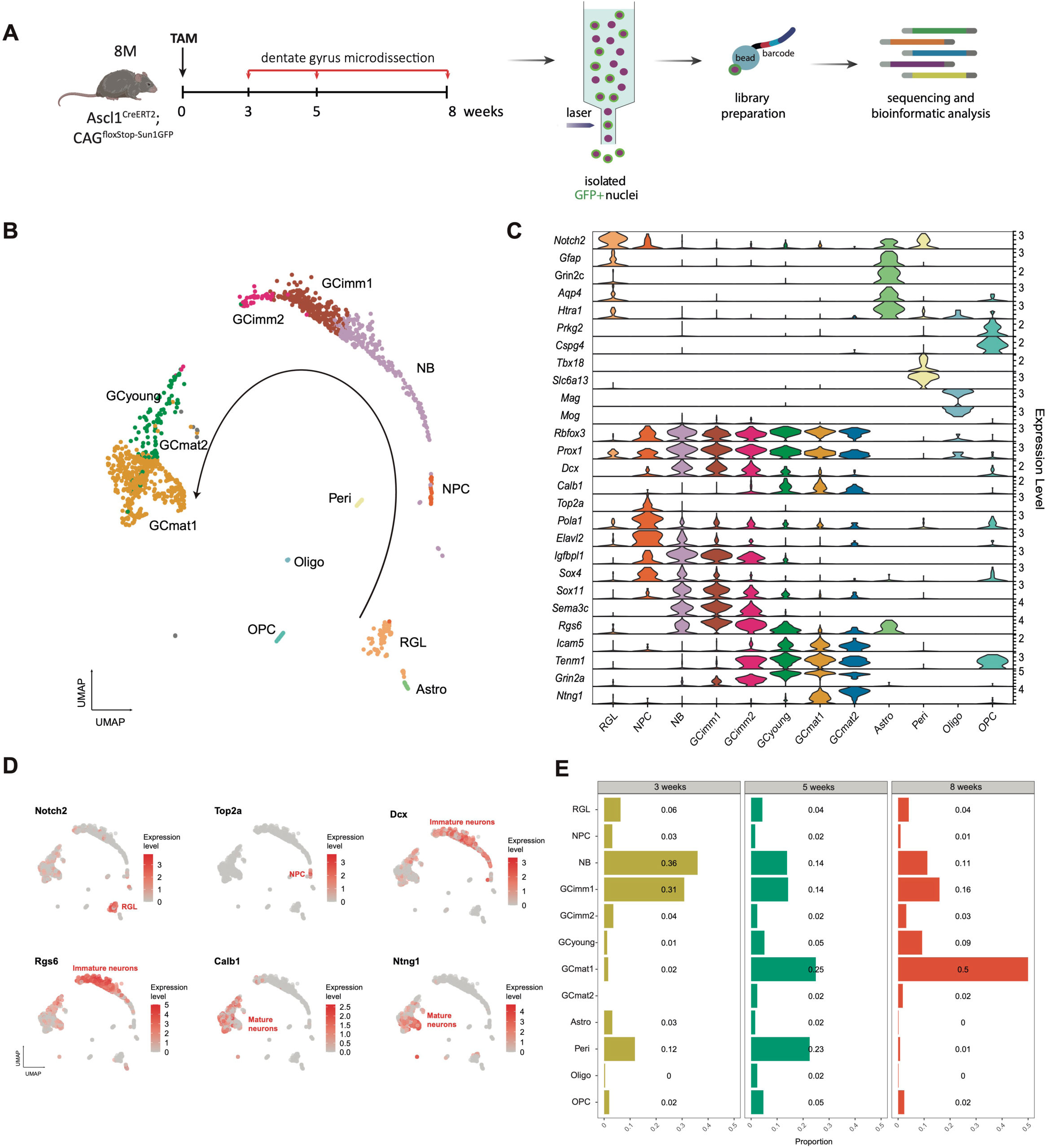
snRNA-seq reveals conserved transcriptional maturation of adult-born dentate granule cells in aging, mirroring young development at a slower pace. (A) Schematic of the experimental workflow. Dentate gyrus tissue was microdissected from *Ascl1^CreERT2^*;*CAG^floxStop-Sun1GFP^* 8M mice at 3, 5, and 8 weeks post-tamoxifen induction. Each cohort of GFP^+^ nuclei was FACS-purified and processed separately for single-nucleus RNA-seq using the 10X Chromium platform. (B) Uniform Manifold Approximation and Projection (UMAP) embedding showing major cell populations identified across developmental stages, including radial glia-like cells (RGL), neural progenitor cells (NPC), neuroblasts (NB), immature, young, and mature granule cells (GCimm1, GCimm2, GCyoung, GCmat1, GCmat2), astrocytes (Astro), oligodendrocytes (Oligo), oligodendrocyte precursor cells (OPC), and pericytes (Peri). Cluster labels were transferred from a previously annotated dataset of two-month-old animals (2M, young dataset) using the Seurat package. Arrow: developmental pathway. (C) Violin plots validating the identified clusters through the expression of canonical marker genes. (D) UMAPs showing the expression of selected canonical marker genes to identify cell populations. *Notch2* (RGL)*, Top2a* (NPCs)*, DCX and Rgs6* (immature neurons), *Ntng1* and *Calb1 (*mature neurons). (E) Proportional representation of each cluster across the 3-, 5-, and 8-week cohorts, showing dynamic changes in cell composition over time.

A birthdating-stamped dataset was generated by combining sorted nuclei with snRNA-seq using 10x-Chromium microfluidics technology. After bioinformatic quality control and filtering, we retained a total of 1925 nuclei with an average of 2036 genes/nuclei, consistent with the expected gene counts for neurons (**Fig. S1A,B**) (26, 29–32). Nuclei were visualized using the UMAP (Uniform Manifold Approximation and Projection) algorithm, which projects them in a low-dimensional phase space. Clusters were identified by transferring labels from a previously annotated dataset obtained from two-month-old mice (2M, young dataset) using the Seurat package (**Fig. 1B**) (26). This approach yielded 12 clusters: oligodendrocyte (Oligo), oligodendrocyte precursor (OPC), pericyte (Peri), astrocyte (Astro), RGL, neural progenitor cell (NPC), neuroblast (NB), immature aGC 1 and 2 (GCimm1 and 2), young aGC (GCyoung), mature aGC 1 and 2 (GCmat1 and 2). Neuronal lineage clusters (from RGL to GCmat) were spatially organized along a linear developmental pathway. The resulting maturation trajectory was consistent with the transcriptional progression occurring during adult hippocampal neurogenesis that we published recently (26). Partition identities were further validated by the expression of canonical markers (**Fig. 1C, D, S1C**).

Neuronal clusters were defined by *Rbfox3*, comprising 7 communities containing 84% of all nuclei. *DCX* (immature neuronal marker) and *Calb1* (mature neuronal marker) allowed the identification of aGCs across different maturation stages (**Fig. 1C, D, S1C**). Neuronal differentiation initiates with NPCs expressing cell cycle genes (*Top2a* and *Pola1),* and NBs expressed *Elavl2* and *Igfbpl1*. Immature neurons were grouped into GCimm1 and GCimm2, defined by *Sox4*, *Sox11*, *Sema3c*, *Rgs6*, and *Dcx*. GCimm2 expressed additional transcripts such as *Tenm1* and *Grin2a*. GCyoung, expressed *Calb1* but lacked *Ntgn1*, which was restricted to the mature granule cell populations, GCmat1 and GCmat2. These partitions were further characterized by expression of their canonical markers: *Icam5, Tenm1* and *Grin2a* **(Fig. 1C)**.

Consistent with our previous young adult mice dataset (26), astrocytes and RGLs were distinguished by the overlapping expression of *GFAP* and *Notch2*. However, *Htra1*, *Grin2c*, and *Aqp4* expression enabled the identification of astrocytes as a distinct cluster from RGLs despite their common lineage (33, 34). Three additional non-neuronal clusters were identified: pericytes expressing *Tbx18* and *Slc6a13*, OPCs expressing *Prkg2* and *Cspg4,* and oligodendrocytes expressing *Mag* and *Mog* **(Fig. 1C, S1C)**.

Our dataset captured the entire trajectory of aGC development in the aging hippocampus. Developmental states were defined on the basis of canonical markers validated in previous studies (26, 31), highlighting the reliability of the transcriptional profiles across ages.

### Temporal dynamics of neurogenesis in the aging brain

A cohort-based analysis revealed key differences in the distribution of cell clusters across developmental timepoints (**Fig. 1E, S1D**). RGLs were detected at low proportions (<10 %) across all cohorts, in accordance with the age-dependent decline in the population of Ascl1-expressing neuronal stem cells (10–12). NPCs were also scarce, reflecting their rapid transition from proliferation to postmitotic maturation states (26, 35). The w3 cohort predominantly contributed to NB and GCimm1, with both populations decreasing at later time points. The proportion of GCimm2 and GCyoung nuclei increased slightly but were scarcely represented at 5 and 8 weeks, suggesting that those cells rapidly progressed through the maturation trajectory. The majority of GCmat1 cells were found at w8, with some contribution from w5. By projecting aged cohort-specific nuclei onto the reference trajectory defined by the young dataset, a subset of nuclei from the w8 cohort remained transcriptionally similar to early developmental stages, suggesting a deceleration in cellular progression **(Fig. S1E)**. These findings indicate that, while the aging neurogenic process retains the cellular identities observed in young mice, it exhibits a marked slowdown in developmental progression, particularly in the young cohort (w3).

This transcriptomic profiling captures different stages of neuronal development in the aging hippocampus, from early differentiation to functional maturation, highlighting a slower developmental trajectory. These observations are consistent with previous morphological and functional studies (9, 17, 18). However, the molecular mechanisms underlying the age-dependent delay remain largely unknown.

### Aging alters transcriptional dynamics and neuronal progression in hippocampal neurogenesis

To uncover the molecular mechanisms underlying the protracted maturation of newborn aGCs in the aging hippocampus, we analyzed the young and aging datasets within a common expression space (**Fig. 2A**). We obtained an overlapped representation that confirmed the alignment between the young-adult and aging partitions. To explore their similarities and differences, we carried out a comparative analysis of differentially expressed genes (DEGs). To minimize bias arising from unequal numbers of nuclei across clusters, the analysis was performed using a bootstrap-based subsampling strategy that repeatedly compared matched numbers of nuclei, retaining only genes consistently identified as differentially expressed in at least 80% of iterations. Interestingly, all clusters revealed similar transcriptional profiles in young adult vs. aging mice, except for RGLs and NBs, which showed a substantial number of downregulated genes, suggesting age-related silencing of specific biological programs (**Fig. 2B; Table S1**). Several genes downregulated in RGLs have previously been implicated in synaptic transmission, synaptic signaling, and neuromodulation, including *Cacna1a, Gabrg3, Gnao1, Plcl1, Grm3, Slc1a3, Plcb1, Kcnq5, Grid2,* and *Prkg1*, suggesting that aging could impair the responsiveness of RGLs to local signaling cues. Similarly, NB downregulated genes were enriched in processes associated with cell maturation, glutamate receptor signaling, and excitability, suggesting potential alterations by aging in programs related to neuronal differentiation (**Fig. S2A; Table S2**).

**Figure 2.**
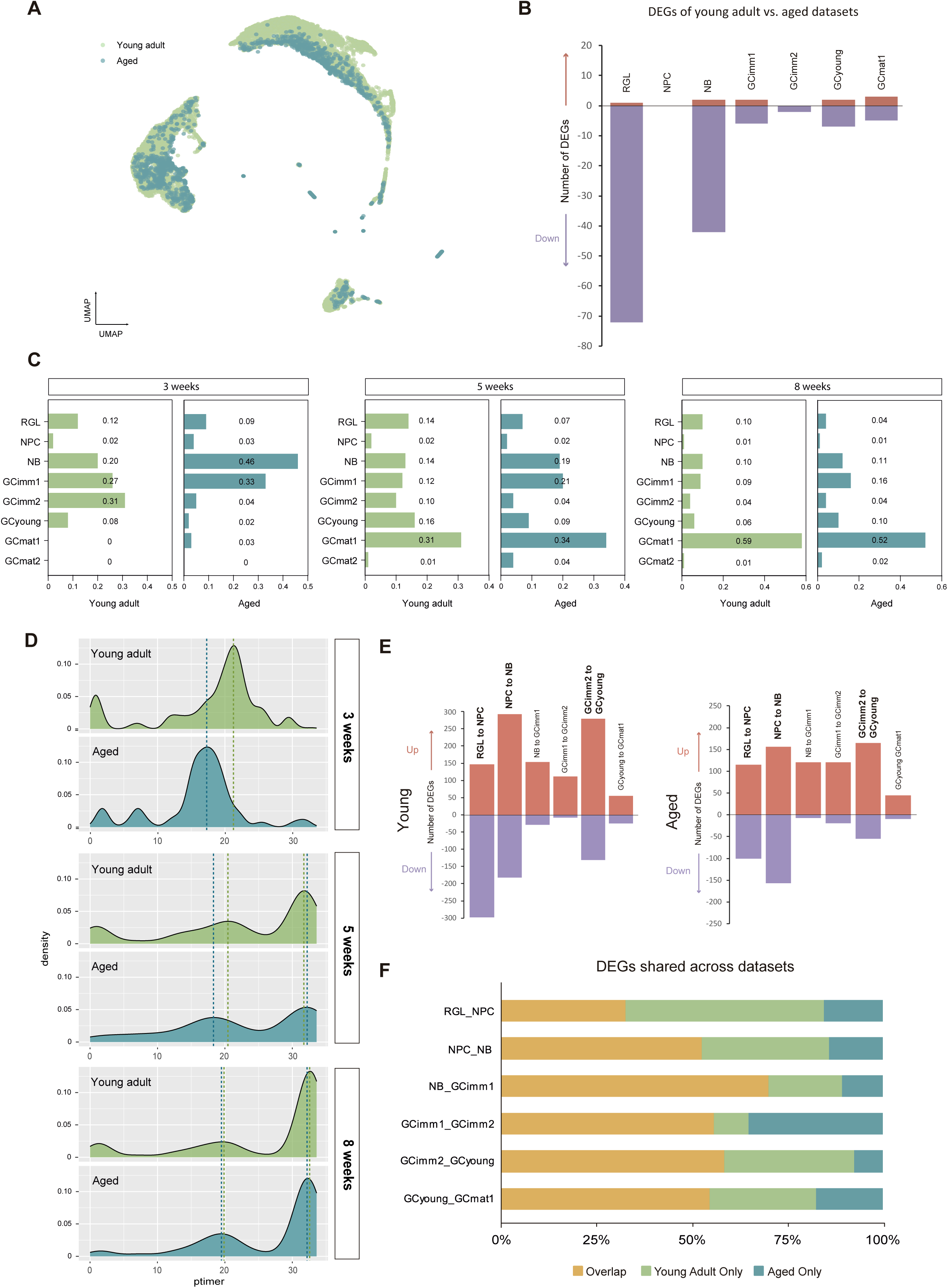
Comparative transcriptional analysis of hippocampal neurogenesis in young adult and aged mice. **(A)** UMAP showing nuclei distribution by age along the first two dimensions. Young adult and aged nuclei are shown in green and blue, respectively. **(B)** Bar plots showing the number of differentially expressed genes (DEGs) between defined clusters for aged vs young adult datasets. Up-and down-regulated genes are shown in pink and violet, respectively (FC ≥ 1.5 or ≤-1.5 and FDR ≤ 0.05). DEGs are listed in Table S1. **(C)** Dynamic changes in cell composition of neuronal clusters across the 3-, 5-, and 8-week cohorts, illustrating proportion for young adult (green) and aged (blue) datasets. **(D)** Density distribution of nuclei along the pseudotime trajectory comparing age-matched cohorts for young adult (green) and aged (blue) datasets. Pseudotime values were assigned based on transcriptomic profiles. Green (young adult) and blue (aged) dashed lines mark density peaks. **(E)** DEGs between adjacent cluster transitions (as indicated above) for young adult (left) and aged (right) datasets. Pink and violet indicate up-and downregulated genes, respectively (FC ≥ 1.5 or ≤-1.5 and FDR ≤ 0.05). DEGs are listed in Table S3. Highlighted transitions represent those with the highest number of DEGs. **(F)** Bar plot showing the percentage of DEGs unique to young (green), aged (blue), or shared (yellow) between datasets during the transition between consecutive clusters.

In addition to these molecular alterations, the proportion of RGLs was low across all cohorts (w3, w5, and w8), consistent with a reduction of the RGL pool in aged animals (**Fig. 2C**). Despite this decline, the overall neurogenic trajectory appeared preserved. However, while developing aGCs reached the GCimm2 stage during the third week (w3) in young adult animals, neurons from middle-aged mice displayed a striking accumulation at the NB stage, suggesting a potential delay in the developmental progression. Moreover, nuclei at the GCimm2 and GCyoung states were scarce, suggesting a low cluster stability along all cohorts in aged mice. The conspicuous developmental delay observed in the w3 cohort was further confirmed by pseudotime analysis (**Fig. 2D**). Density plots revealed shifts in the timing of peak cell densities, particularly a delayed progression for the w3 cohort in aged mice. However, positions of peak cell densities became progressively aligned in w5 and w8 cohorts.

While cluster identities remained similar, the altered dynamics in the stage progression in the aging dataset suggested possible differences in gene regulatory networks. To better understand the transcriptional complexity underlying these changes, we compared DEGs across successive clusters within the young and aging datasets **(Fig. 2E; Table S3)**. The overall DEG pattern shared features across ages. Transitions from RGL to NPC, NPC to NB, and GCimm2 to GCyoung displayed high numbers of downregulated genes, consistent with key developmental shifts from quiescence to a proliferative program (RGL to NPC), from proliferation to a postmitotic state (NPC to NB), and from immature to fully mature aGCs (GCimm2 to GCyoung) (26). In contrast, the NB to GCimm2 transitions were dominated by transcriptional activation, with few genes being silenced (**Fig. 2E; Table S3**). Upregulated genes included synaptic organizers and adhesion molecules (*Dlg2, Lrfn5, Ncam2, Cntnap2, Cntn1, Cadm1, Lrrtm4, Kirrel3, Cdh9, Cdh11, Cdh12*), neurotransmission components (*Grin2a, Grm1, Grm5, Grm7, Gria3, Gabbr2, Gabrb1, Gabrb3*), axon guidance and neurite outgrowth regulators (*Sema6d, Sema3c, Sema5a, Epha5, Epha6, Epha7, Robo1, Dcc, Dscam*), and cytoskeletal modulators (*Prickle2, Carmil1, Nav2, Nav3, Shtn1, Map1b*). Notably, all of these upregulated genes are common to the young adult and aging datasets. These transcriptional changes reflect an intense phase of synaptic specification, connectivity refinement, and structural maturation. Conversely, downregulated genes comprised additional adhesion molecules and synaptic regulators (*Lrrc4c, Lrrtm3, Ctnna3, Tenm4*), transcriptional and developmental regulators (*Sox5, Maml3, Msi2, Zmiz1*), and signaling modulators (*Ppp2r2b, Pik3r1, Pde4b, Plcb1, Hcn1*), suggesting the repression of early developmental and plasticity-associated programs in favor of a functionally committed aGC identity.

Interestingly, the proportion of DEGs shared between datasets was high across most transitions (>50%), with the notable exception of RGL to NPC (**Fig. 2F**). We thus focused on this transition, which exhibited a markedly reduced number of downregulated genes in aging mice compared to the young dataset (**Fig. 2E, S2B**). Downregulated genes that were restricted to the young dataset revealed significant associations with Wnt signaling (*Tle4, Ctnnbip1, Znrf3, Gli3, Prkn, Tcf7l2, Prickle1, Foxo1, Nxn, Eda*) and stem cell proliferation (*Gli3, Fgfr1, Id4, Ptch1, Znrf3*) (**Table S3**). Transcripts also included regulators of canonical Wnt/β-catenin signaling (e.g., *Ctnnbip1, Tcf7l2, Znrf3*) or noncanonical Wnt pathways (e.g., *Nxn*), both of which are essential for adult hippocampal neurogenesis (36–38). The lack of downregulation of these genes in aging mice suggests an alteration in the transcriptional switch that normally accompanies the transition from activation to quiescence. These differences may impair the proper regulation of RGL proliferation and contribute to the reduced neurogenic potential observed in the aging brain.

### Aging aGCs exhibit developmental stalling at the neuroblast state

The RNA-seq analysis shows results consistent with the delayed neurogenesis in the aging DG (9). To validate the different transcriptomic dynamics across ages, we performed RNAscope *in situ* hybridization analysis on brain tissue sections to visualize early stages of the developmental trajectory of aGCs. Specific marker combinations were used to distinguish NPCs (*Elavl2^+^*), NBs (*Elavl2^+^* / *Sema3C^+^*), and GCimms (*Sema3C^+^*) (**Fig. 3A**). Imaging via deconvolution microscopy enabled precise detection of single and co-expressed transcripts within individual cells (**Fig. 3B-D**). Young adult mice displayed primarily NBs and GCimms, while DGs from aging mice showed a larger population of NBs but a diminished proportion of GCimms (**Fig. 3E; Table S4**). It is striking that immature GCs are the most abundant population of developing neurons in the young adult dentate gyrus, while the aging hippocampus is dominated by NBs. This is consistent with the neuroblast accumulation and delayed progression toward immature granule cell states revealed by transcriptomic analysis.

**Figure 3.**
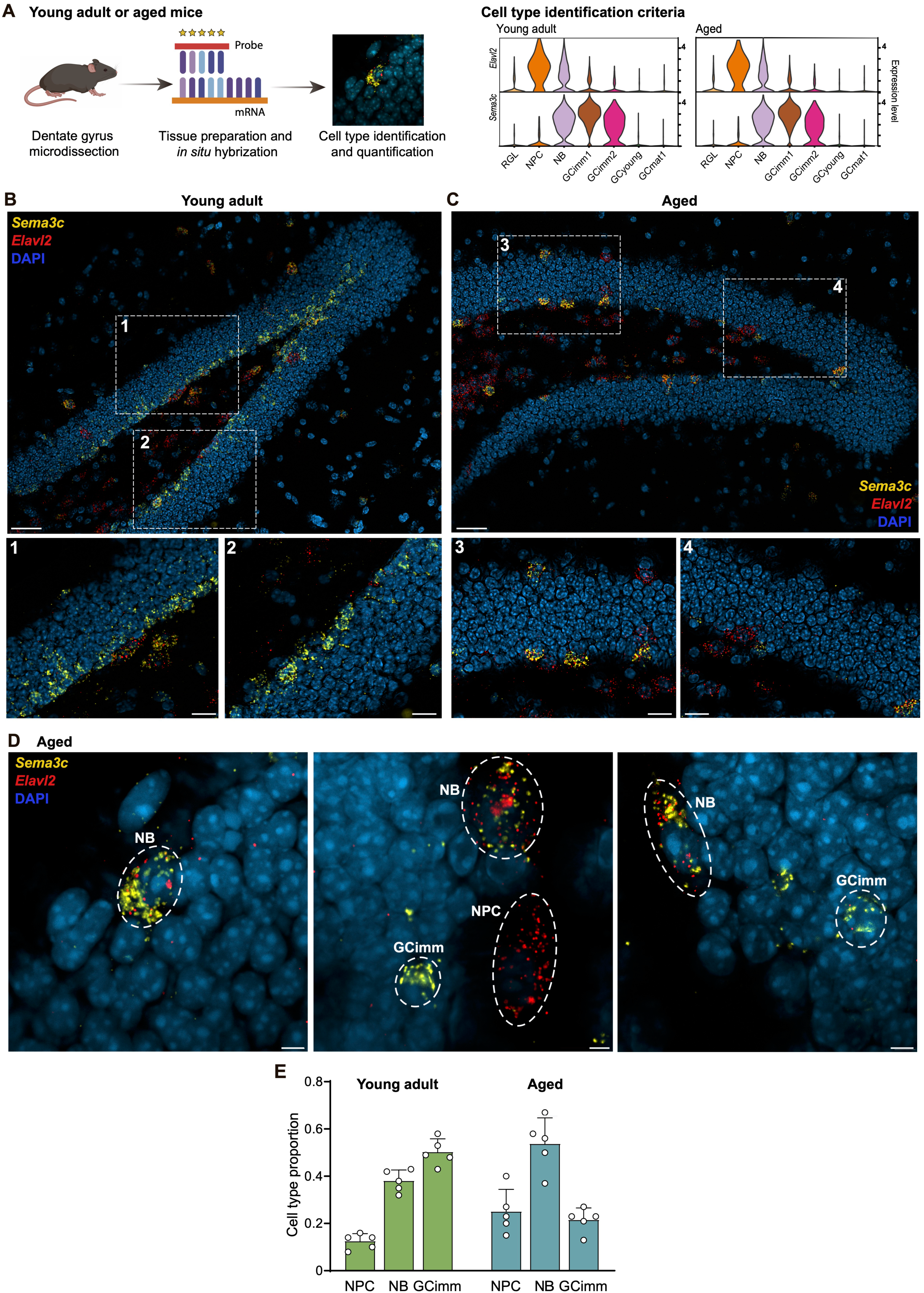
Aging impairs neuronal maturation by stalling adult-born granule cells at the neuroblast stage. **(A)** Schematic of the RNAscope strategy used to identify distinct neuronal populations in the dentate gyrus of young adult and aged mice. Neural progenitor cells (NPCs) were defined as *Elavl2*⁺, neuroblasts (NBs) as *Elavl2*⁺/*Sema3C*⁺, and immature granule cells (GCimms) as *Sema3C*⁺. Given that both *Elavl2* and *Sema3C* are also expressed in non-neuronal cell populations, only marker-positive cells located within the SGZ were included in quantitative analyses. **(B, C)** Representative deconvoluted images showing the spatial distribution of labeled cells in the dentate gyrus of young adult (**B**) and aged mice (**C**). Panoramic views (top panels; scale bar: 50 μm) and magnified insets (bottom panels; scale bar: 20 μm) illustrate reduced neurogenesis in aged brains, along with increased *Elavl2/Sema3C* co-localization, indicating NB accumulation. DAPI (blue) marks nuclei. **(D)** High-resolution Airyscan images showing single-labeled and co-labeled cells, enabling unambiguous classification of NPCs, NBs, and GCimms (scale bar: 5 μm). **(E)** Quantification of cell populations reveals a shift toward increased NBs and reduced immature granule cells in aged mice, consistent with a developmental stalling at the NB stage. In contrast, young adult animals exhibit a progressive increase in GCimms (N = 5 animals per group). Data are shown as means ± SEM. Cell counts are listed in Table S4.

### Neuroblasts are the primary neuronal state modulated by physical exercise

The accumulation of cells at the NB stage may represent a developmental bottleneck in the aged hippocampus, reflecting impaired progression toward neuronal maturation. Previous studies have demonstrated that voluntary running accelerates adult neuronal differentiation in the young and aged brain (9, 18, 39). To investigate the developmental transitions and critical molecular pathways modulated by exercise, we compared 3-week-old aGCs of aged mice from sedentary or voluntary running conditions throughout the entire developmental period (**Fig. 4A**). Using the same pipeline as before (**Fig. 1A**), we projected nuclei together with the young dataset into a shared expression space (**Fig. 4B**). This analysis showed that cells from the running group were shifted towards more mature clusters compared to sedentary controls. Furthermore, analysis of the distribution of cell cohorts along the different clusters revealed that running mice exhibited >50% reduction in NBs, accompanied by increased proportions of GCimm2 and GCyoung (**Fig. 4C**). To further evaluate the dynamics of neuronal progression under different conditions, we performed a comparative pseudotime analysis (**Fig. 4D**). Nuclei from sedentary aged mice displayed a dominant peak that corresponded to cells within early immature differentiation. The aged running group displayed a bimodal distribution, with a dominant peak reaching a position closer to the young cohort, and an additional peak corresponding to more advanced developmental stages. These findings demonstrate that running promotes neuronal maturation in the aging hippocampus by releasing developing aGCs from a stalled NB stage.

**Figure 4.**
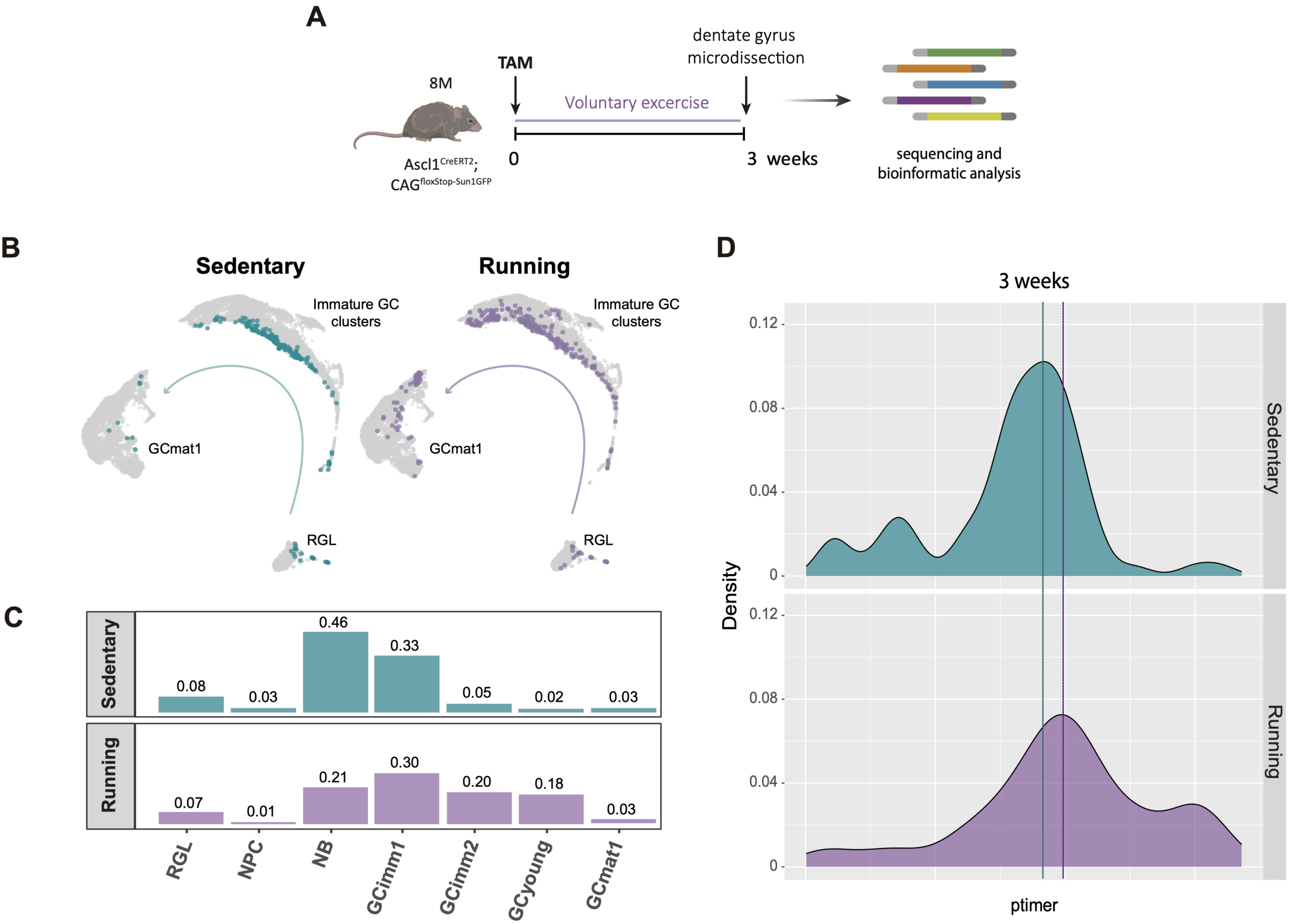
Voluntary exercise promotes the transition from neuroblasts to immature and young granule cells in the aging dentate gyrus. (A) Schematic of the experimental design. Aged *Ascl1^CreERT2^*;*CAG^floxStop-Sun1GFP^* mice received tamoxifen (TAM) injections and were assigned to voluntary running or sedentary housing for three weeks, followed by dentate gyrus microdissection. GFP⁺ nuclei were FACS-purified and processed separately for single-nucleus RNA-seq using 10x Chromium technology. (B) UMAP embeddings of cells from sedentary (blue) and running (violet) aged mice, illustrating temporal progression along the developmental trajectory derived from the young dataset. (C) Proportions of cell clusters in sedentary (blue) and running (violet) aged mice, showing a ∼50% reduction in NBs and increased GCimm and GCyoung in the running group. **(D)** Density distribution of nuclei along the pseudotime trajectory in aged sedentary (top, blue) and aged running (bottom, purple). Pseudotime values were assigned based on transcriptomic profiles. Dashed lines indicate density peaks.

### Molecular changes underlying exercise-induced neuronal maturation

To investigate how voluntary running influences the molecular maturation of new aGCs, we analyzed DEGs in w3 cohorts from aged sedentary and running mice (**Fig. 5A; Table S5**). Upregulated genes in the running group were significantly enriched for GO terms linked to neuronal maturation and connectivity, including glutamate receptor signaling, synapse assembly, and axon guidance, consistent with an enhanced structural and functional integration (**Fig. 5B**). Interestingly, w3 neurons from runners exhibited transcriptional profiles closely resembling those of w5 aGCs from sedentary mice (**Fig. 5C**). To further highlight this shift, we focused on the 34 transcripts showing the strongest upregulation (FC>1.8) in runners. These genes, encompassing synaptic organizers (*Lrrtm4, Pcdh9, Ncam2, Lrfn5*), axon guidance cues (*Epha5, Epha6, Sema6d, Lingo2*), and regulators of neuronal excitability and synaptic plasticity (*Kalrn, Gabrb1, Kcnd2, Fgf14*), displayed expression levels in neurons from aging runners that were positioned between w5 and w8, rather than w3 cells in sedentary mice (**Fig. 5D)**. The expression of these genes was found to be particularly elevated in the mature neuronal clusters of young mice (**Fig. S3A**). Together, these findings suggest that voluntary exercise accelerates the molecular trajectory of newborn aGCs, promoting the early acquisition of transcriptional programs characteristic of more mature developmental stages.

**Figure 5.**
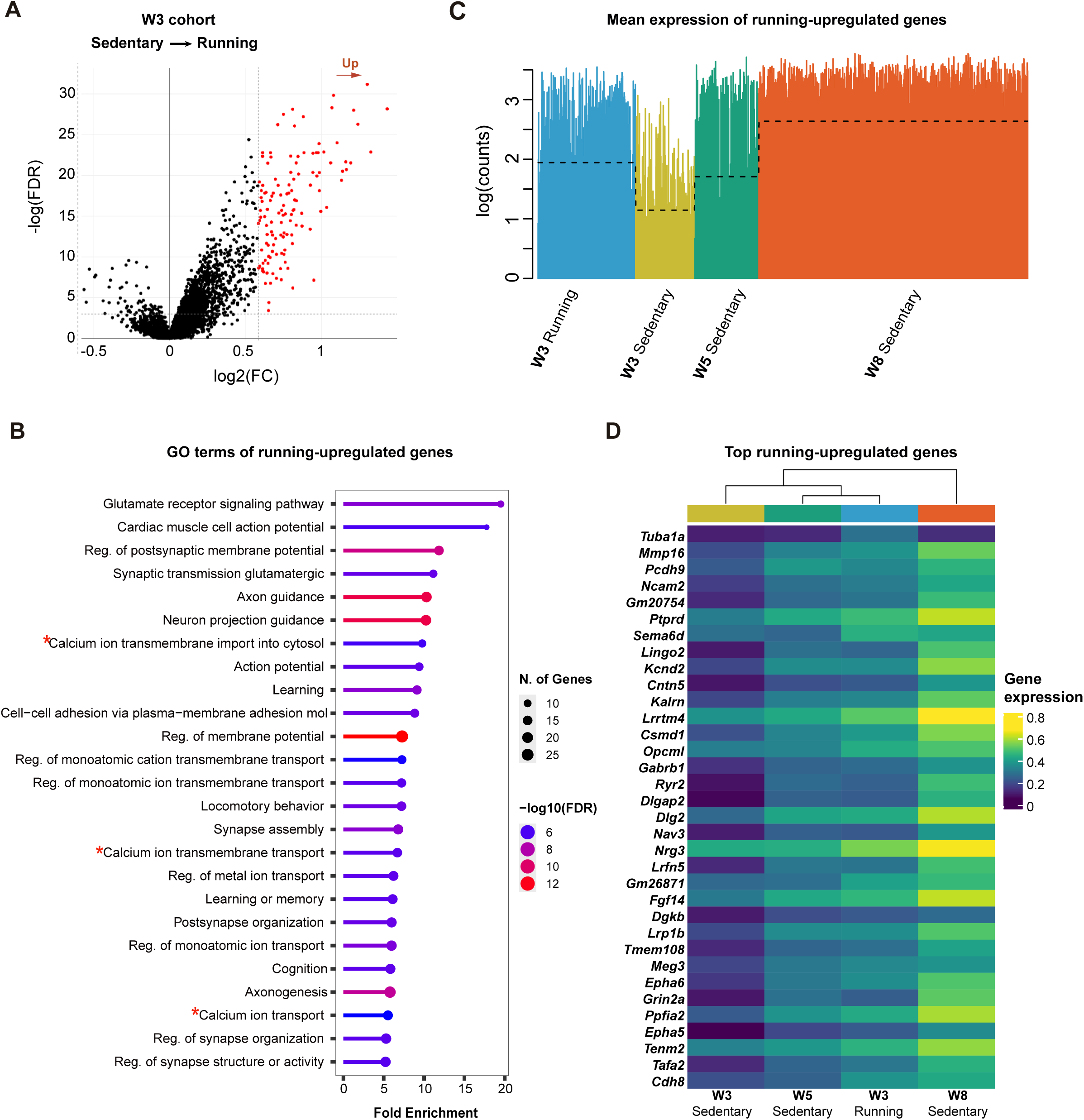
Voluntary running induces transcriptional changes associated with structural and synaptic reorganization. **(A)** Volcano plot showing differential expression analysis (FC ≥ 1.5 or ≤-1.5 and FDR ≤ 0.05) of 3-week-old aGCss from sedentary versus running mice. DEGs are listed in Table S5 **(B)** GO enrichment analysis of upregulated genes in running aged mice, with processes related to synaptic signaling and organization, structural axonal and dendritic remodeling, and cognitive functions. Enrichment analysis was performed using ShinyGO v0.85, with a false discovery rate (FDR) cutoff of 0.05. Pathways are sorted according to - Log10(FDR). GO are listed in Table S5 **(C)** Spike plots displaying mean expression of 127 upregulated genes of 3-week-old GCss from sedentary versus running aged mice (FC ≥ 1.5 or ≤-1.5 and FDR ≤ 0.05). Black dashes show the mean expression for each cohort. **(D)** Heatmap of hierarchical clustering for the expression of 34 upregulated genes with highest fold-change (FC>1.8) from sedentary versus running aged mice. Scale on the right denotes mean expression level.

To determine whether these transcriptional changes were shared across the entire population or restricted to specific cell states, we analyzed differentially expressed genes (DEGs) within the most abundant clusters in our samples (**Fig. S3B; Table S5**). NB, Gcimm1 and GCimm2 displayed minimal differences between conditions, supporting the notion that running accelerated the trajectory rather than producing within cluster modifications.

Notably, the extent of transcriptional change observed in the entire 3-week-old neuronal population (127 upregulated genes in runners) exceeded that seen within all single clusters together (**Fig. 5A, S3A**). The bulk cohort of 3-week-old neurons encompasses a heterogeneous mix of maturation states. In sedentary mice, this population is enriched in NBs and GCimms, whereas in runners, it is predominantly composed of more developmentally advanced GCimm2 neurons (**Fig. 4C**). This shift in cell-type proportions likely contributes to the greater transcriptomic divergence observed at the population level. Together, these findings reinforce the postmitotic NB stage as a key checkpoint in adult neurogenesis during aging that remains responsive to activity-dependent cues.

### Running reduces apoptosis during early stages of aGC maturation

The transient delay in aGC maturation observed in the w3 cohort of aged mice was rescued by voluntary running, and gradually recovered in later sedentary cohorts (**Fig. 4, 2C**). Although initially interpreted as a developmental delay, accumulating evidence suggests that aging neuroblasts exhibit reduced survival capacity, raising the possibility that the w3 bottleneck reflects an impaired survival in addition to the delayed differentiation (40).

To determine whether this developmental delay involves altered survival, we examined apoptotic activity in immature aGCs from sedentary and running aged mice (**Fig. 6A**). As DCX is expressed by both NBs and GCimms, active Casp3/DCX immunostaining was used to assess apoptosis in these early neuronal stages. Sedentary aged mice displayed a larger proportion of DCX⁺/Casp3⁺ cells than running animals, suggesting that physical activity prevents apoptosis and facilitates the transitions towards more advanced stages (**Fig. 6B,C; Table S4**).

**Figure 6.**
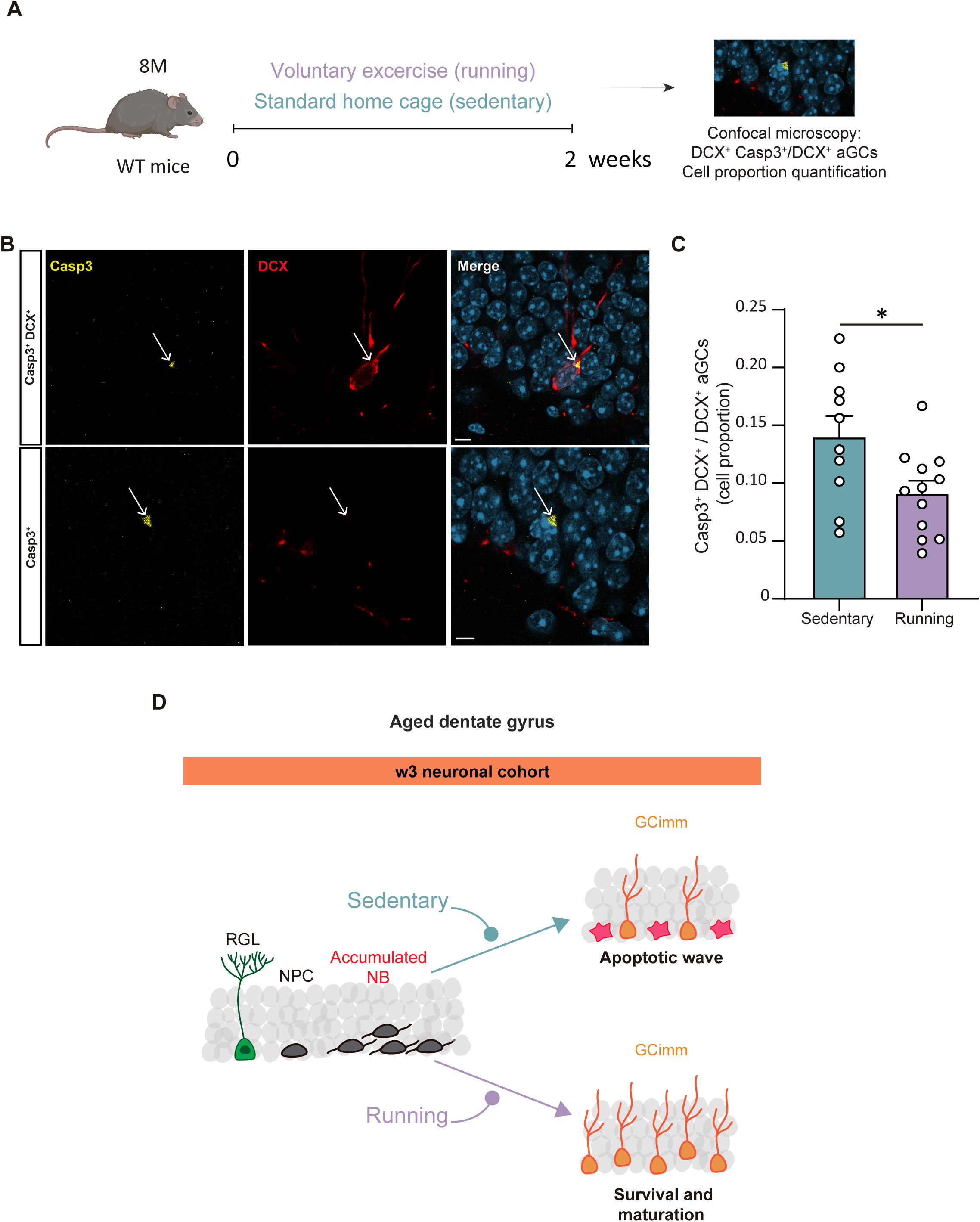
Running reduces apoptosis during early stages of aGC maturation in the aged dentate gyrus. **(A)** Scheme of the experimental design. Aged WT mice were assigned to voluntary running or sedentary housing for two weeks, followed by confocal imaging to count Casp3⁺DCX⁺ cells, normalized to the total number of DCX⁺ cells per dentate gyrus. **(B)** Representative confocal images showing active Casp3⁺DCX⁺ colocalized cells (upper panel) and active Casp3⁺DCX^-^ cells (lower panel) in the dentate gyrus of running mice. Scale bars: upper panel, 10 µm; lower panel, 5 µm **(C)** Quantification of Casp3⁺DCX⁺/DCX⁺ cells shows a significant decrease in apoptosis in running mice compared with sedentary controls (unpaired *t-test*, *P* = 0.039; each dot represents one DG; N = 5 mice (sedentary), 6 mice (running), 2 DG per mouse). Data are shown as means ± SEM. Cell counts are listed in Table S4. **(D)** *Effect of physical activity on the fate of adult-born neurons in the aged dentate gyrus.* Schematic representation of a 3-week-old neuronal cohort in sedentary (top) and running (bottom) aged mice. NPCs generate NBs that frequently accumulate in aged conditions. Under sedentary conditions, most newborn cells undergo an apoptotic wave, resulting in a reduced number of NBs differentiating into GCimm. In contrast, physical activity alleviates this bottleneck by promoting NB survival and progression, enabling their maturation into GCimm and GCyoung. RGL: radial glia-like cell; NPC: neuronal progenitor cell; NB: neuroblast; GCimm: immature granule cell.

Because the running condition upregulated GO categories related to calcium import into the cytosol (**Fig. 5B**), and activity-dependent Ca²⁺ signaling is known to promote neuronal survival during hippocampal development, we next examined transcriptional markers associated with Ca²⁺ entry and downstream pro-survival pathways (41, 42). Notably, the L-type Ca²⁺ channel subunits *Cacna1c* and *Cacna1d*, which mediate depolarization-evoked Ca²⁺ influx, as well as *Akt3*, a key survival kinase, were upregulated in running animals (**Fig. S3C**). Given that L-type calcium channel activity and Akt signaling are essential components of the PI3K/Akt survival pathway in developing hippocampal neurons, these data support the idea that running enhances Ca²⁺-dependent pro-survival signaling during early stages of aGC maturation in the aged DG.

Together, these findings suggest that the NB bottleneck detected in the w3 cohort represents a survival-dependent checkpoint, at which NBs stall while awaiting survival or activity-dependent cues. In this context, physical activity supports not only the maturation pace and overall neurogenesis rate, but also preserves early aGCs survival, allowing stalled neuroblasts to resume maturation in the aged DG (**Fig. 6D**).

## DISCUSSION

Recent advances in single-cell and spatial transcriptomics have provided unprecedented insight into the aged hippocampus, identifying molecular signatures associated with aging and more importantly, the beneficial effects of physical exercise (24, 25). Yet, despite these advances, the transcriptional dynamics underlying adult hippocampal neurogenesis during aging have remained largely unexplored. By combining genetic lineage tracing techniques with cell-type label transfer from young reference datasets, we demonstrate that transcriptional identities of aGC developmental stages are largely preserved in the aging hippocampus, but their population dynamics differ (26). Anchoring cell identities across young and aged samples ensured that the observed differences reflect changes in neurogenic dynamics rather than shifts in cluster definition. This strategy revealed age-related alterations in the composition and transcriptional trajectories of neurogenic cohorts that would be difficult to interpret using *de novo* clustering approaches alone. Although this strategy constrains the discovery of age-specific clusters, it provides a robust framework to interrogate how aging reshapes the dynamics of adult neurogenesis within conserved cellular states.

Overall, the transcriptional identities of aGC developmental stages were largely conserved with age, and the extent of differential gene expression across transitions along the differentiation trajectory was remarkably similar between young and aging brains (**Fig. 2E**). The main exception emerged at the RGL-to-NPC transition, where the number of downregulated genes in the aged brain was half the value detected in the young brain. Consistent with this observation, the RGL-to-NPC transition displayed the lowest overlap in shared DEGs (**Fig. 2F**), further supporting that RGL states diverge substantially between the young and aging hippocampus. Accordingly, we detected the most pronounced transcriptional differences across datasets at the RGL stage, consistent with recent evidence indicating that aging alters the regulatory landscape governing stem cell quiescence and activation in the hippocampus (13, 43). In line with this interpretation, RGLs were underrepresented (<10%) in the more advanced cohorts compared with young datasets (**Fig. 2C**), likely reflecting both a reduction in Ascl1-expressing cells and a decreased recruitment of stem cells into the RGL-activated pool (11, 44–46).

Beyond this difference in the RGL cluster, the most striking change along the neurogenic trajectory was the accumulation of postmitotic NBs within the w3 cohort, accompanied by a delayed transition towards more mature GC stages. The convergence of lineage-resolved transcriptomic profiling, RNAscope validation, and apoptosis analyses supports the interpretation that aging is associated with an accumulation of neurons at the NB stage accompanied by increased vulnerability to cell death. These findings align with previous observations of protracted morphological maturation in aGCs from aged animals, and provides the molecular resolution to pinpoint the discrete stage at which developmental progression becomes constrained (9, 17, 18).

Previous morphological and electrophysiological studies have demonstrated that voluntary running can rescue age-associated delays in aGC maturation (9, 17, 18). Our data refine this view by showing that running accelerates the progression of newborn neurons along a preserved differentiation trajectory, thereby promoting faster maturation (**Fig. 5, S3**). In aged running mice, the w3 cohort shifted toward a distribution dominated by GCimm2 neurons (**Fig. 4C**). GCimm2 cells express transcriptional signatures characteristic of functional neurons capable of establishing synaptic connections (26). These findings complement previous reports showing that running profoundly reshapes the molecular landscape of the hippocampal niche beyond aGCs, influencing astrocytes, microglia, and oligodendrocyte precursor cells (24). Such widespread remodeling of the neurogenic niche may contribute to the activity-dependent signals that facilitate neuronal maturation in the aging hippocampus. Our findings suggest that the period of increased apoptotic vulnerability during aging neurogenesis presented at the NB stage can be partially alleviated by voluntary running. In line with previous results, the aged brain retains a substantial neurogenic potential, and external cues, likely relayed through the neurogenic niche, play a predominant role in unlocking this latent capacity.

Despite the pronounced redistribution of cell states induced by running, only modest transcriptional differences were detected within the NB cluster, which might reflect a transient nature of activity-dependent transcriptional responses. Previous work has shown that behavioral stimulation rapidly induces transcriptional programs in aGCs that emerge within hours and subsequently return to baseline (47). Thus, many of the acute transcriptional responses might no longer be detectable after prolonged voluntary running. In this context, the primary consequence of running may be dominated by the altered cell-state composition and developmental progression rather than persistent transcriptional changes.

The identification of this bottleneck raised the possibility that survival is compromised during early maturation stages. Indeed, we observed a marked apoptosis among DCX⁺ aGCs in aged sedentary mice, reflected by the proportion of Casp3⁺/DCX⁺ cells, which was substantially reduced in runner mice. This apoptotic wave matches the developmental window during which a large fraction of newborn neurons is normally eliminated, a process that appears exacerbated in aging (48). Remarkably, running significantly reduced Casp3⁺ NBs, demonstrating that physical activity enhances early-stage neuron survival. Together, these findings identify a reversible apoptotic checkpoint at the NB stage that contributes to age-related decline in neurogenesis. The mechanisms underlying this checkpoint likely reflect a convergence of immune, metabolic, and activity-dependent signals, acting both at the cell intrinsic and extrinsic levels. For instance, aged NBs may express senescence-associated markers, rendering them vulnerable to immune-mediated elimination (40). At the same time, NBs can undergo apoptosis if they fail to receive adequate trophic signals or synaptic inputs during the early postmitotic period (48). Our data further show that running upregulates transcriptional programs related to Ca²⁺ import and neuronal maturational pathways. Therefore, physical activity may counteract NBs apoptotic vulnerability by increasing trophic and activity-dependent signaling.

Taken together, these observations support the role of early postmitotic NBs as a critical regulatory node where aging disrupts neuronal progression. External interventions such as physical exercise can relieve this constraint by promoting maturation and enhancing survival. We propose a model in which the aged dentate gyrus retains a reservoir of newborn neurons that accumulate at the NB stage and remain responsive to pro-maturational cues. This accumulation may reflect a prolonged window during which neuronal survival and maturation remain sensitive to activity-dependent signals. A conceptually related phenomenon was recently reported using 40 Hz audiovisual stimulation, which enhances the rapid integration of new aGCs in the aging brain (49). These findings support the notion that in the aging dentate gyrus distinct forms of network activation can recruit stalled neuronal cohorts into functional circuits. From this perspective, this aging-associated accumulation at the neuroblast stage, although it limits neuronal progression, also leaves a pool of immature neurons poised for activity-dependent maturation.

## ACKNOWLEDGMENTS

We thank Ignacio Satorre for technical assistance, and members of the A.F.S., A.C., and Emilio Kropff labs for insightful discussions. We are grateful to the Microscopy and Bioimaging Center (CeMBio) at the Leloir Institute for assistance with confocal imaging. D.G., A.C., M.F.T. and A.F.S. are investigators in the Consejo Nacional de Investigaciones Cientificas y Tecnicas (CONICET). N.B.R, M.H., and A.A.B. were supported by CONICET fellowships. P.A. serves as a Scientific Advisory Board (SAB) member at Foresite Labs and CNSII, and is a cofounder and SAB member of Vesalius Therapeutics. This work was supported by NIH grants from the National Institute of Neurological Disorders and Stroke (NINDS) R01NS128117 (PA), NINDS and Fogarty International Center R01NS103758 (AFS and PA), and the Argentine Agency for the Promotion of Science and Technology PICT 2017-0389 (DG), PICT 2018-03713 (AC), and PICT-2020-0046 and PICT-2021-0077 (AFS).

## MATERIALS AND METHODS

### Animals

Male and female C57Bl/6J wild type and genetically modified mice (8-9 months of age) were housed at four to five mice per cage in standard conditions. For adult neurogenesis snRNA-seq experiments, *Ascl1^CreERT2^ (Ascl1^tm1(Cre/ERT2)Jejo/J)^*) mice were crossed to *CAG^floxStop-Sun1/sfGFP^* (B6.129*-Gt(ROSA)26Sort^m5.1(CAG-Sun1/sfGFP)Nat/MmbeJ^*) conditional reporter line to generate Ascl1^CreERT2^;CAG^floxStop-Sun1/sfGFP^ mice, which were used to reliably target adult-born GC nuclei (28, 50, 51). Tamoxifen (TAM) induction (120 µg/g, three injections in three consecutive days) resulted in the expression of Sun1-sfGFP in the nuclear envelope of Ascl1^+^ cell progeny. Mice were anesthetized (150 μg ketamine/15 μg xylazine in 10 μl saline per g) and sacrificed at the indicated times after TAM induction: 3-, 5-and 8 weeks. All animals were maintained in C57BL/6J background. Experimental protocols were approved by the Institutional Animal Care and Use Committee of the Leloir Institute (CICUAL-FIL 85), according to the Principles for Biomedical Research involving animals of the Council for International Organizations for Medical Sciences and provisions stated in the Guide for the Care and Use of Laboratory Animals. Leloir Institute is approved as a foreign facility by the Office of Laboratory Animal Welfare of the US National Institutes of Health (F18-00411).

### Tissue dissection

Mice were deeply anesthetized (ketamine/xylazine, as described above), brains were carefully removed and placed into ice cold Earl’s balanced salt solution (EBSS, in mM: 117 NaCl, 5.4 KCl, 1 NaH_2_PO_4_, 26 NaHCO_3_, 5.6 glucose, 1.8 CaCl_2_.2H_2_O and 0.8 MgSO_4_) with trehalose 5% v/v and kynurenic acid 0.8 mM (52). Dissection solution was equilibrated in 95% O_2_/5% CO_2_ before use. Dentate gyrus was dissected and processed following a previous established protocol (26). The dissected tissue was placed in a 0.5 ml microcentrifuge tube with a minimal amount of media, flash-freezed on dry ice and stored at-80°C until use.

### Nuclei isolation and FACS sorting

Nuclei were isolated as previously described with several modifications (30). Dounce tissue grinder and pestles were sequentially washed with 100% EtOH, RNAse Zap (Sigma, Cat # R2020), 2-3 rounds with RNAse-free water, and finally rinsed with EZ Lysis Buffer (Sigma, Cat # NUC-101). All material and buffers were chilled on ice. The tissue was transferred to a chilled dounce prefilled with 2 ml ice-cold EZ Lysis Buffer and homogenized slowly with 10 strokes of pestle A, followed by 10 strokes with pestle B. Suspension was transferred to a chilled tube and incubated on ice 5 min. Then the suspension was centrifuged at 500 × g for 5 min at 4°C and the supernatant was removed. The isolated nuclei were resuspended in 4 ml ice-cold Nuclei Suspension Buffer (NSB) (1x RNase-free molecular-biology grade PBS, 0.1% molecular-biology grade BSA 100 μg/ml and 0.2U/μl RNase inhibitor). Finally, nuclei were centrifuged at 500 × g for 5 min at 4°C and after removing the supernatant were resuspended in 1ml NSB-Ruby (final Ruby concentration 1:500) and filtered twice through a 35-μm cell strainer to remove as many debris as possible. Nuclei were kept on ice until sorting (Harvard University, Bauer Core Facility, BD FACSAria II) into 96 well plates, pre-coated with BSA to reduce adherence and improve recovery, containing with ∼10-20 μl of rich-NSB (with final 2U/μl of RNase inhibitor and 1% BSA). The final nuclei concentration was determined using a C-chip Neubauer counting chamber and Trypan Blue (1:2 final dilution). Nuclei were immediately loaded for single-cell GEM formation (10x Genomics, single cell RNA sequencing 3ʹ, Chromium v3.1).

### 10x Genomics Chromium

Dataset sampling for snRNA-seq experiments was carried out within a minimal time interval, processing two time points in a single experiment each day to minimize batch effects. For each time point, four dentate gyri from male and female mice were pooled. For snRNA-seq, nuclei were loaded in the 10x Genomics chips aiming to recover 2,000–10,000 nuclei. cDNA amplification and library construction were done following 10x Genomics protocols. For both dataset 1 and dataset 2, libraries were generated using Chromium v3.1, quantified in BioAnalyzer and sequenced on an Illumina platform. Samples were sequenced to a depth of 40–70,000 reads per cell.

### RNA *in situ* hybridization

For cell type identification, 5 young and 5 aged WT mice were anesthetized and the brains were removed and placed into the ice-cold dissection solution (described above). Hippocampi were dissected under a microscope and immediately embedded in OCT on dry ice and stored at-80 °C. Sections of 14 μm covering the anterior–posterior axis of the dentate gyrus were collected in a cryostat. Sections were placed for 1 h at-20 °C and stored at-80 °C until use. RNA *in situ* hybridization was performed using the RNAscope Fluorescent Multiplex Reagent Kit (Advanced Cell Diagnostics), according to the manufacturer’s instructions. Briefly, thawed sections were fixed in PFA 4%, dehydrated in sequential incubations with ethanol, followed by 30 min protease IV treatment. Appropriate hybridization probes (Advanced Cell Diagnostics, catalogue#: *Sema3c* 441441-C1, *Elavl2* 491961-C2,) were incubated for 2 h at 40°C, followed by amplification steps (according to protocol), DAPI counterstaining, and tissue was mounted with gerbatol to prevent bleaching.

### Immunofluorescence

Immunostaining was performed in 60 μm-thick free-floating coronal sections throughout the septal fraction of the hippocampus from retrovirally injected mice. Antibodies were applied in TBS with 3% donkey serum and 0.25% Triton X-100. Immunofluorescence was performed using the following primary antibodies: DCX (Goat, 1:250, Santa Cruz #8066), Caspase-3 (Rabbit, 1:250, Cell Signalling #9664). The following corresponding secondary antibodies were used: donkey anti-rabbit Cy5 and donkey anti-goat Cy3 (1:250, Jackson ImmunoResearch Laboratories). Incubation with DAPI (10 minutes) for nuclear counterstain was performed. Slices were mounted and covered with gerbatol to prevent bleaching.

### Microscopy

Sections from the septal hippocampus (−0.94 to −2.46 mm from bregma, according to the mouse brain atlas (53) were analyzed. For experiments shown in Figs. 3B and 3C, images were acquired using deconvolution software (Zen Blue 3.3; Zeiss) with a 20×/0.8 NA objective. Airyscan images (**Fig. 3D**) were obtained using an LSM 880 Airyscan microscope (Zeiss) with a 63×/1.4 NA objective from 14-µm-thick sections. Confocal images (**Fig. 6B**) were acquired on the same system using a 63×/1.4 NA from 60-µm-thick sections.

### snRNA-seq processing

We used STARsolo 2.7 to align the snRNA-seq reads to the GRCm38 mouse genome. Multi-mapped reads were excluded and both exonic and intronic counts were considered. We used default parameters to count UMI and filter high-quality cells in order to generate gene-by-cell count matrices. Quality control (QC) and downstream data processing were performed using *scran* and *scDblFinder* bioconductor libraries (54, 55). All scripts used to perform the present analysis were included in the companion website https://github.com/chernolabs/NeuronalSwitch. We eliminated 353 low-quality nuclei out of 2613 nuclei that passed the STARsolo quality filter. Low-quality nuclei were identified by either a multivariate outlier detection procedure (*adjOutlyingness* function of robustbase R library) based on low library sizes, low number of detected features and large mitochondrial (Mt) content, or by presenting more than 1% of Mt counts. Features with >20 molecules and detected in >1% and <90% of filtered nuclei were retained. In addition, we removed specific genes (*Ehd2, Espl1, Jarid1d, Pnpla4, Rps4y1, Xist, Tsix, Eif2s3y, Ddx3y, Uty, Kdm5d, Rpl26, Gstp1, Rpl35a, Erh, Slc25a5, Pgk1, Eno1, Tubb2a, Emc4, Scg5*) to mitigate possible sex and stress effects on downstream analysis (31). The data was normalized using the scran library in accordance with the OSCA pipeline (56). We executed a quick clustering for each timepoint independently, normalized nuclei in each cluster separately, and rescaled the size factors to be comparable across clusters. Finally, we computed the log2 of the data and considered a pseudo-count value of 1 to generate a normalized expression matrix. After scaling and log-normalizing gene expression values, we used the scran *modelGeneVar* function to identify the 3000 most variable genes in their log-expression profiles.

Using the interactive *shiny* interface provided by the **scX** R package (https://chernolabs.github.io/scX/), we visually inspected the dataset and detected nuclei expressing marker genes corresponding to choroid plexus cells (*Ttr, Htr2c, Enpp2*) and interneurons (*Gad1* and *Gad2*). These nuclei were classified as non–dentate gyrus contaminants and were excluded from further analyses (262 nuclei).

We used Seurat’s label transfer to annotate nuclei based on a juvenile dentate gyrus reference dataset (26). Two nuclei were labeled as NB2 and subsequently removed from the analysis. We determined differentially expressed genes between nuclei groups using the *FindMarkers* function from the *scran* library (*t*-test pairwise). Cluster marker genes were identified as highly ranked differentially expressed genes between the query group and all other clusters.

### Analysis of differential gene expression

DEG analysis in Fig. 2B was performed by subsampling nuclei to avoid bias due to unequal sample size across clusters. A bootstrap approach was applied, and only genes identified as differentially expressed in at least 80% of iterations were retained.

### Pseudotime determination

The *slingshot* 2.4.0 R-package was used to fit developmental trajectories to nuclei of the following clusters: RGL, NPC, NB1, GCimm1, GCimm2, GCyoung and GCmat1 (57). We considered the UMAP (3D) low dimensional coordinates to fit principal curves and produce pseudotime estimations.

### Visualization and graphical reports

We used Force Atlas 2 layout algorithm, implemented in Gephi to produce 2-dim visualization of mKNN graphs (58). Heatmaps, volcano plots, violin plots, dot plots, and spike plots were produced using the scX R package [https://chernolabs.github.io/scX/] (59).

### Gene Ontology analysis

GO enrichment analysis was performed using ShinyGO v0.85. The analysis was conducted applying a false discovery rate (FDR) cutoff of 0.05. Only GO terms associated with a minimum of 20 genes and a maximum of 500 genes were considered (http://bioinformatics.sdstate.edu/go/) (60).

### Statistical analysis

All data are presented as mean ± SEM. Analyses were performed blind to the operator. Immunofluorescence colocalization and quantification were verified using Zen Blue 3.2 (Zeiss). Statistical analyses were conducted using GraphPad Prism 8.0.2 (GraphPad Software, CA, USA). Two-group comparisons were performed using unpaired Student’s *t*-test after confirming normality with Anderson-Darling test, D’Agostino–Pearson, Shapiro–Wilk, and Kolmogorov–Smirnov tests.

## SUPPLEMENTARY FIGURES

**Figure S1. Quality control, cluster distribution and canonical marker gene expression. (A)** Summary of experimental data showing post-tamoxifen timepoints (3, 5, and 8 weeks), number and sex of mice (F: female, M: male), number of FACS-sorted and analyzed nuclei (#: number). **(B)** Quality control of snRNA-seq libraries. Scatter plots show the number of detected genes (features) versus the number of mRNA molecules (counts) per nucleus for each timepoint, indicating comparable sequencing depth across cohorts. **(C)** UMAP showing expression of canonical marker genes used to identify neuronal and non-neuronal populations. Markers include *Htra1* (pericytes), *Cspg4* (OPCs), *Slc6a13* (astrocytes), *Mog* (oligodendrocytes), and *Rbfox3* (neurons), supporting accurate cluster annotation. **(D)** Distribution of nuclei across clusters for each neuronal cohort. **(E)** UMAP embedding for individual aged cohorts, projected along the reference trajectory defined by the young dataset (Rasetto et al, 2024), illustrate temporal progression along the developmental path. Nuclei from the young dataset are shown in gray along the maturation trajectory.

**Figure S2. Gene ontology enrichment and age-specific transcriptional changes during adult neurogenesis. (A)** Top enriched GO pathways among genes downregulated in aged vs. young NBs. GO and genes are listed in Table S2. **(B)** Proportion of downregulated (left) and upregulated (right) genes unique to young (green), aged (blue), or shared (yellow) during the transition between consecutive clusters.

**Figure S3. Effect of physical activity on the number and expression of genes. (A)** UMAP visualization for the expression levels of the 34 most upregulated genes in running vs. sedentary cohorts at 3 weeks. **(B)** Bar plots showing the number of DEGs between defined clusters for sedentary and running mice (w3 cohort for both of the conditions). Upregulated (pink) and downregulated (violet) genes are indicated (FC ≥ 1.5 or ≤-1.5 and FDR ≤ 0.05). DEGs are listed in Table S5. **(C)** Dot plot showing the normalized expression for *Cacna1c, Cacna1d* and *Akt3* for the 3-week cohort from sedentary versus running aged mice. Scale on the right denotes mean expression level.

## SUPPLEMENTARY TABLES

**Table S1. Related to Figure 2B.** DEGs identified across clusters in the young adult versus aged datasets. Columns include gene name, direction of regulation (Up, upregulated; Down, downregulated), and number of occurrences across 100 iterations.

**Table S2. Related to Fig. S2A.** GO enrichment analysis of genes downregulated in the comparison of young adult versus aged NBs (Fig. S2B).

**Table S3. Related to Fig. 2E.** DEG analysis across consecutive clusters (transitions) in the young adult and aged datasets, including transitions from RGL to NPC, NPC to NB, NB to GCimm1, GCimm1 to GCimm2, GCimm2 to GCyoung, and GCyoung to GCmat.

**Table S4. Related to Fig. 3E and 6C.** Cell counts used for quantification. For Fig. 3E, columns include the number of quantified NPC, NB, and GCimm cells in young-adult and aged mice. For Fig. 6C, counts include the number of DCX^+^ neurons, as well as DCX^+^ and Casp3^+^ cells per dentate gyrus in aged mice under sedentary and running conditions.

**Table S5. Related to Fig. 5A, B, and S3B.** DEG analysis in nuclei from 3-week-old cells in aged mice comparing sedentary versus running conditions (Fig. 5A). GO enrichment analysis of genes upregulated in sedentary versus running comparisons (Fig. 5B). DEG analysis within the same clusters in sedentary versus running conditions (Fig. S3B).

