## Supplemental Figures 1-3 for "Cell-state mapping reveals a reversible neuroblast accumulation in the aging mouse hippocampus"

### Figure S1

**A**

| Time points | #mice | sex | # FACS events | # nuclei |
|---|---|---|---|---|
| w3 | 4 | F, M | 503 | 240 |
| w5 | 4 | F, M | 1035 | 262 |
| w8 | 4 | F, M | 2150 | 1096 |
| w3 run | 4 | F, M | 245 | 398 |
| **Total** | | | | **1996** |

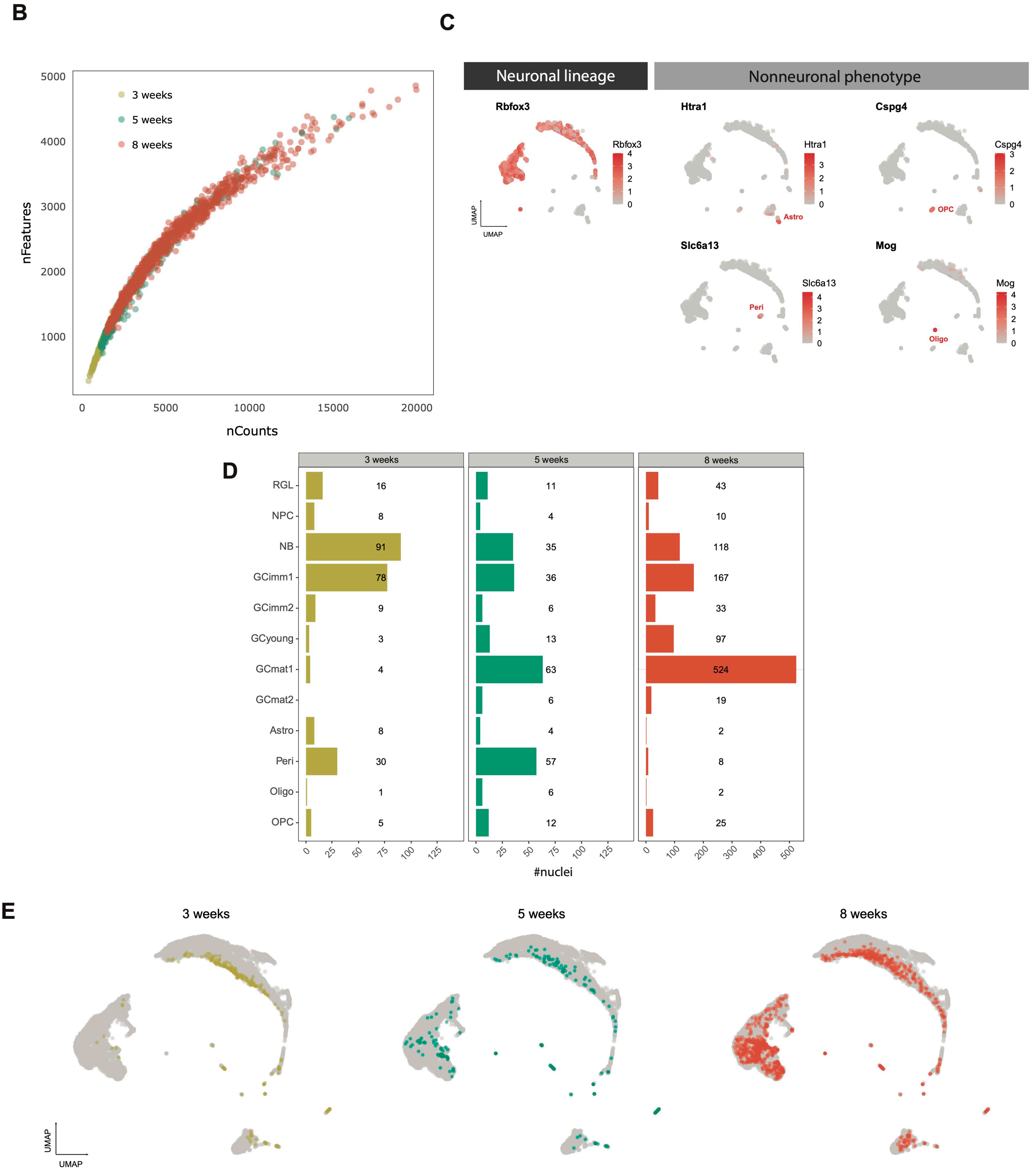

### Figure S2

**A**

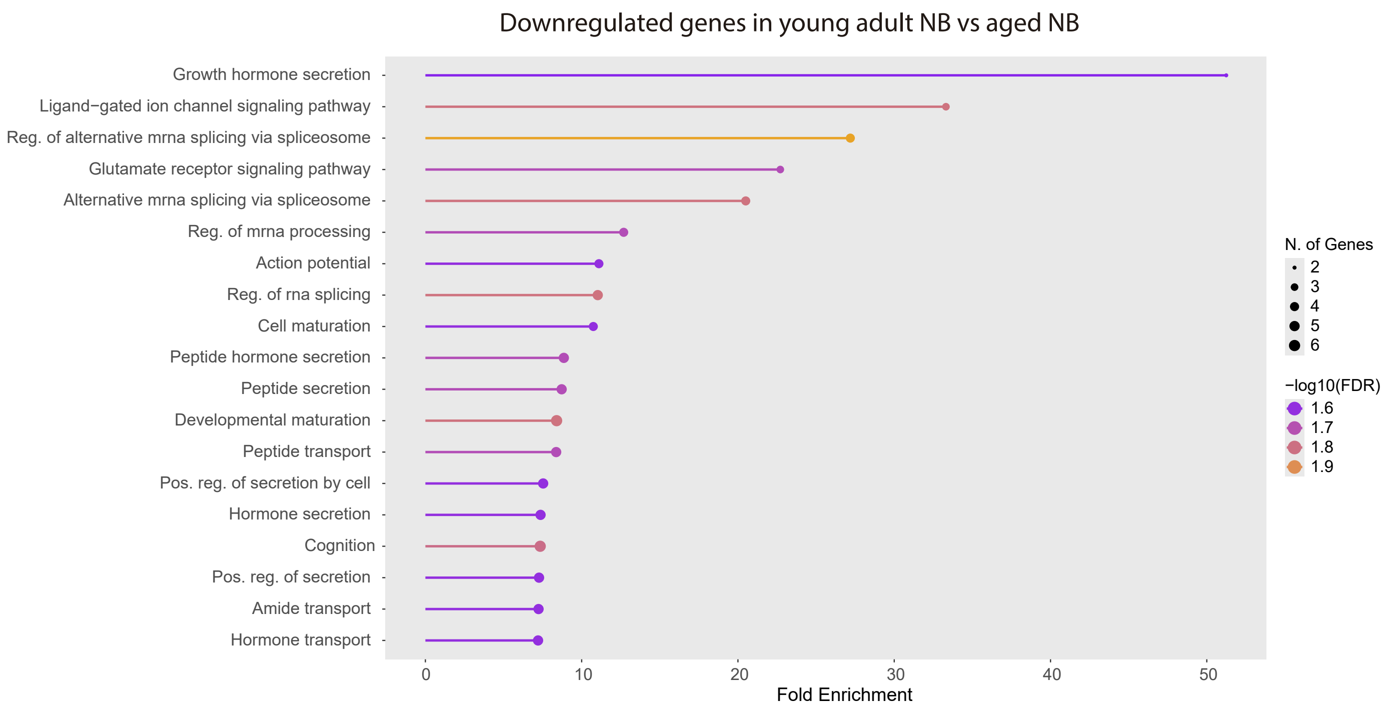

Downregulated genes in young adult NB vs aged NB
Growth hormone secretion
Ligand−gated ion channel signaling pathway
Reg. of alternative mrna splicing via spliceosome
Glutamate receptor signaling pathway
Alternative mrna splicing via spliceosome
Reg. of mrna processing
Action potential
Reg. of rna splicing
Cell maturation
Peptide hormone secretion
Peptide secretion
Developmental maturation
Peptide transport
Pos. reg. of secretion by cell
Hormone secretion
Cognition
Pos. reg. of secretion
Amide transport
Hormone transport
0
10
20
30
40
50
Fold Enrichment
N. of Genes
2
3
4
5
6
−log10(FDR)
1.6
1.7
1.8
1.9


**B**

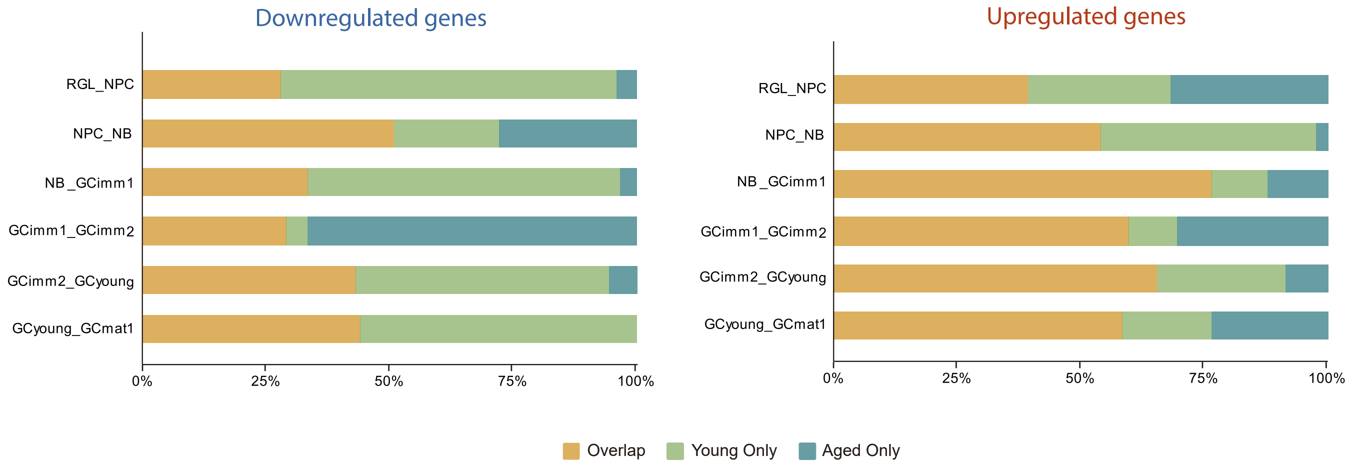

Downregulated genes
Upregulated genes
RGL_NPC
NPC_NB
NB_GCimm1
GCimm1_GCimm2
GCimm2_GCyoung
GCyoung_GCmat1
0%
25%
50%
75%
100%
Overlap
Young Only
Aged Only

### Figure S3

A

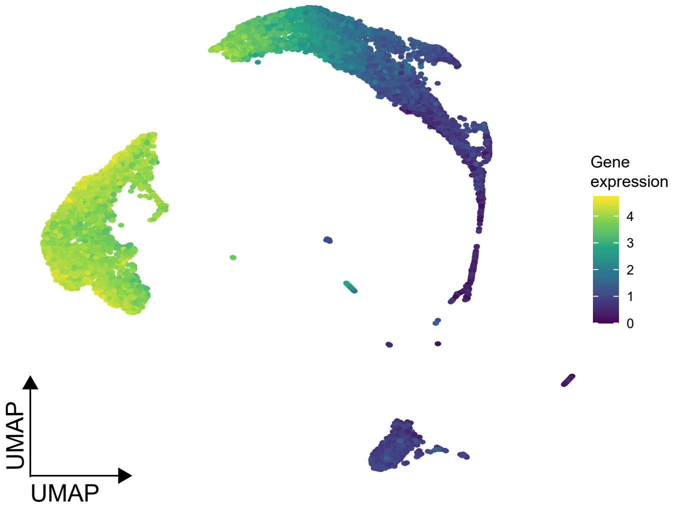


B

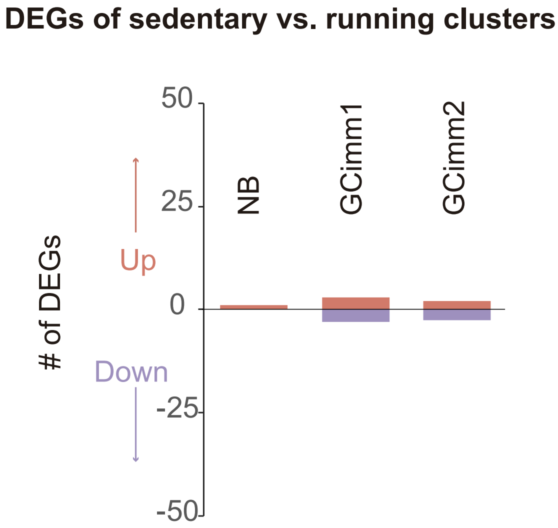


C

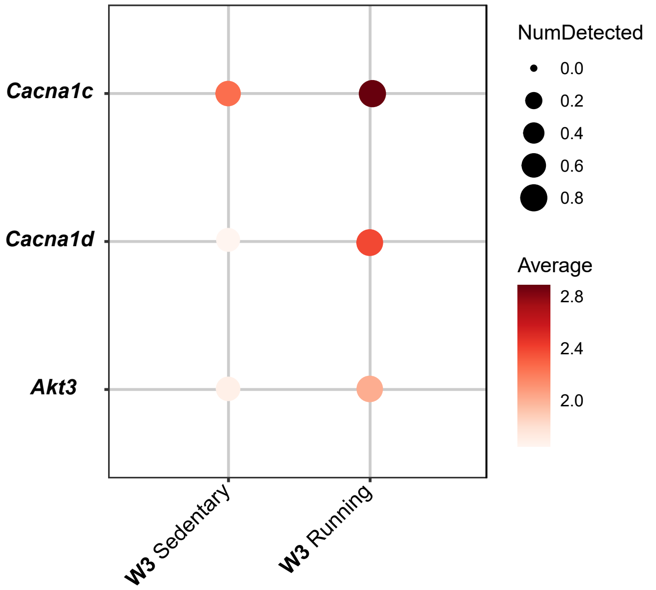
